# Enrichment of non-B-form DNA at *D. melanogaster* centromeres

**DOI:** 10.1101/2021.12.20.473213

**Authors:** Venkata S. P. Patchigolla, Barbara G. Mellone

## Abstract

Centromeres are essential chromosomal regions that mediate the accurate inheritance of genetic information during eukaryotic cell division. Despite their conserved function, centromeres do not contain conserved DNA sequences and are instead epigenetically marked by the presence of the centromere-specific histone H3 variant CENP-A (centromeric protein A). The functional contribution of centromeric DNA sequences to centromere identity remains elusive. Previous work found that dyad symmetries with a propensity to adopt non-canonical secondary DNA structures are enriched at the centromeres of several species. These findings lead to the proposal that such non-canonical DNA secondary structures may contribute to centromere specification. Here, we analyze the predicted secondary structures of the recently identified centromere DNA sequences from *Drosophila melanogaster*. Although dyad symmetries are only enriched on the Y centromere, we find that other types of non-canonical DNA structures, including DNA melting and G-quadruplexes, are common features of all *D. melanogaster* centromeres. Our work is consistent with previous models suggesting that non-canonical DNA secondary structures may be conserved features of centromeres with possible implications for centromere specification.

## Introduction

Eukaryotes share a common mechanism to faithfully segregate genetic information during each cell cycle by which chromosomes are attached to microtubule fibers and are physically pulled towards opposite poles by the kinetochore. Centromeres are essential chromosomal regions that specify the site for the assembly of the kinetochore and are epigenetically marked by chromatin enriched in the histone H3 variant centromeric protein A (CENP-A). CENP-A has been shown to be sufficient for kinetochore assembly and *de novo* recruitment of CENP-A in *D. melanogaster* somatic cells (Chen et al., 2014; Mendiburo et al., 2011; Palladino et al., 2020). Despite their conserved and essential function, centromeres are among the most rapidly evolving regions of genomes (Melters et al., 2013). This rapid evolution has been proposed to be a result of intra-genomic conflict whereby centromeres act as selfish genetic elements driving the rapid evolution of centromeric proteins (Henikoff et al., 2001; Malik and Henikoff, 2009). Furthermore, in organisms such as fungi, nematodes, insects, plants, and vertebrates, centromere function is largely independent of the presence of centromeric DNA sequences, relying instead on the presence of CENP-A chromatin (reviewed in (McKinley and Cheeseman, 2016)). Thus, for most species, the functional significance of centromeric DNA sequences in dictating (or at least contributing to) centromere identity remains unclear.

In an effort to identify genetic characteristics shared amongst the centromeres of diverse eukaryotes, Kasinathan et al. (Kasinathan and Henikoff, 2017) surveyed centromeric DNA sequences from mouse, chicken, *S. pombe* and humans for the presence of <10-bp dyad symmetries (a.k.a. inverted repeats), which are known to adopt unconventional secondary structures such as stem-loops or cruciform extrusions. The authors found that the centromeres of species such as the African Green monkey, chicken, and the fission yeast *S. pombe* were enriched in these motifs. Centromeres enriched in dyad symmetries also showed a predicted propensity to form non-canonical secondary DNA structure under stress, such as that resulting from DNA supercoiling caused by transcription or replication. Non-canonical DNA structures are known as non-B-form DNA and collectively represent any deviation from double stranded B-DNA (the right-handed helix with 10-nt per turn). High likelihood of predicted cruciforms correlated with enrichment in dyad symmetries and other structures, such as melt DNA, were also predicted for some species. Interestingly, centromeres devoid of dyad symmetries, such as those of humans, contain binding sites for CENP-B, a protein that binds specifically to CENP-B box DNA motifs found within α-satellite (Verdaasdonk and Bloom, 2011). CENP-B binding results in the bending of DNA (Tanaka et al., 2001), which in itself represents another non-canonical DNA structure. Based on these analyses, the authors proposed that non-canonical secondary structures may have been selected for during centromere evolution, with a possible role as a structural cue for centromere specification (Kasinathan and Henikoff, 2017). Various non-B structures such as hairpins (Jonstrup et al., 2008), R-loops (Kabeche et al., 2018) and i-motifs (Garavis et al., 2015a; Garavis et al., 2015b) have been observed *in vitro* and *in vivo*, consistent with this model. How widespread centromeric non-B-DNA structures across species may be remains unknown.

The centromeres of *D. melanogaster* were not identified and characterized until recently through a combination of long-read sequencing, chromatin immunoprecipitation, and OligoPaints Fluorescence In-Situ Hybridization (FISH). Chang et al. identified five contigs that make up at least part of the centromeres (Chang et al., 2019) (**Fig. 1A**). The contigs for centromeres X, 3 and 4 contain an island of complex DNA enriched in retroelements flanked by simple satellite repeats. For centromere 2, only a short contig was identified, which contains a small island with a single truncated retroelement flaked by simple satellites. Lastly, the contig for the Y centromere consists of a large island and no satellite DNA. FISH on mitotic chromosomes and extended chromatin fibers show that for centromeres X, 2 and 4, the CENP-A domain spans a region larger than the contig itself, which, based on cytological analyses, can be inferred to be made up of unassembled simple satellites (Chang et al., 2019).

**Figure 1:**
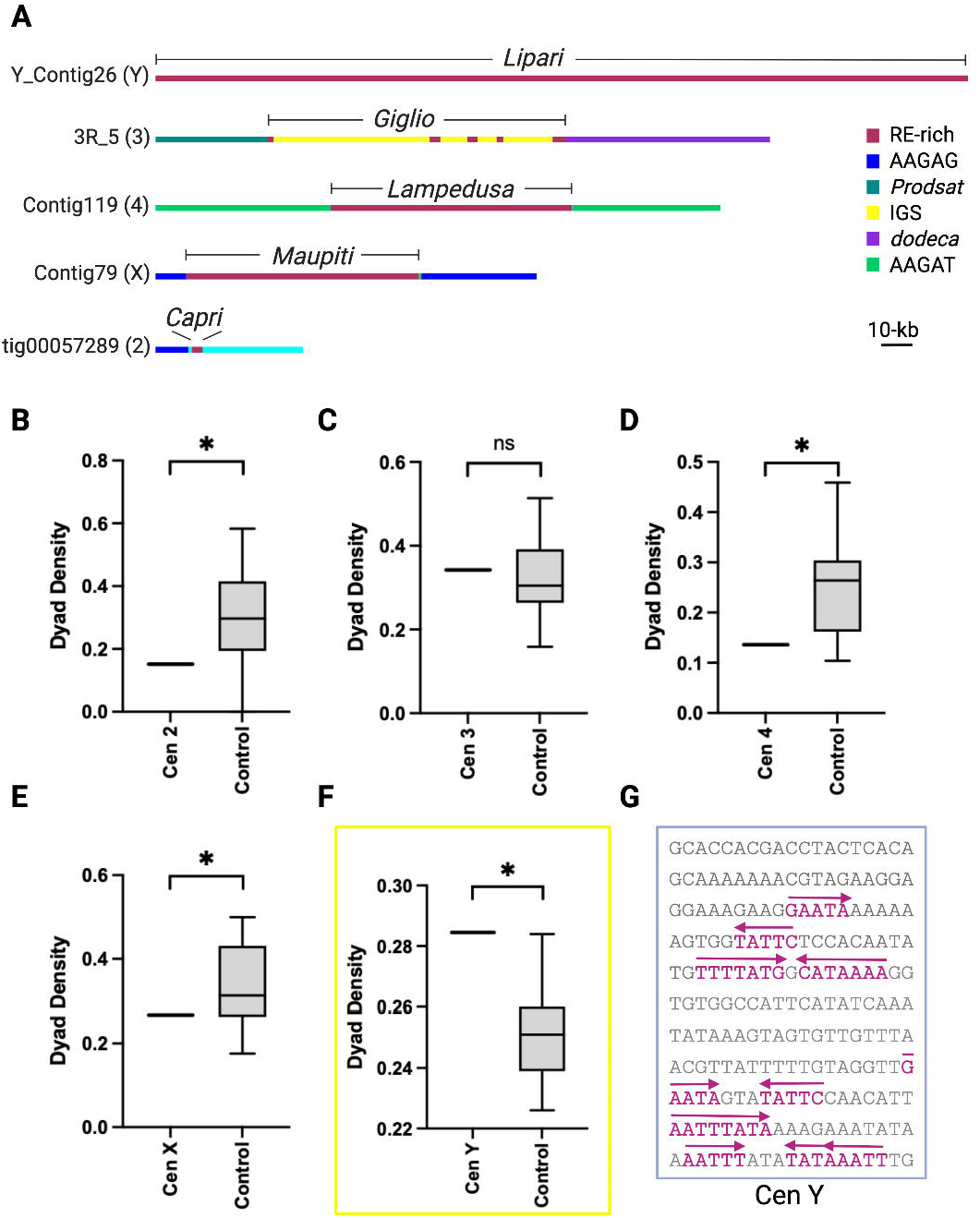
Dyad symmetries are not common features of *D. melanogaster* centromeres. (A) Schematic of the DNA organization of *D. melanogaster* centromere contigs. (B-F) Dyad symmetry density plots for *D. melanogaster* centromeres. Only the Y contig (Y_Contig26; yellow box) showed a significant enrichment. P<0.05, one-sample t-test. (G) Example of inverted repeats from the Y centromere contig (base pairs 181–390).

Here, we use several prediction algorithms to survey the presence of non-B-DNA-form at the centromeres of *D. melanogaster*. Although we show that inverted repeats and cruciform extrusions are not a predominant feature at *D. melanogaster* centromeres, we find evidence for the enrichment of other predicted non-canonical secondary structures such as melted DNA and G-quadruplexes.

## Results and discussion

### Dyad Symmetries are not common features of *D. melanogaster* centromeres

To determine if *D. melanogaster* centromeres are enriched in <10-bp DNA dyad symmetries as previously reported for the centromeres of other species (Kasinathan and Henikoff, 2017), we used the program Palindrome from the EMBOSS suite. We used five contigs (one for each of the X, 2, 3, 4 and Y chromosomes) that are highly enriched in CENP-A chromatin immunoprecipitations and were confirmed to be associated with CENP-A using OligoPaint FISH on extended chromatin fibers as the bona fide *D. melanogaster* centromeres (Chang et al., 2019) (**Fig. 1A**). For our controls, we used several composition and length-matched random genomic sequences for each of the centromere contigs (see Methods). We plotted the EMBOSS palindrome output by calculating the dyad density, obtained by adding the number of base pairs that are part of a dyad divided by the sequence length, and found that only the Y centromere displays dyad symmetry densities higher than control average (**Fig. 1B-G**). These analyses suggest that dyad symmetries are not major features of *D. melanogaster* centromeres and thus are unlikely to play a role in centromere specification. A lack of dyad symmetries was previously reported for human, great apes and *M. musculus* centromeres (Kasinathan and Henikoff, 2017).

### Enrichment of predicted non-B-form DNA structures at centromeric contigs using SIST

The EMBOSS palindrome algorithm identifies dyad symmetries based on sequence analysis. However, this algorithm does not take into account the predicted thermodynamics of DNA and thus does not provide information on the secondary structures it is likely to adopt. Superhelical transitions occur in DNA when negative supercoiling drives susceptible regions to acquire forms alternative to native B-DNA that are energetically favorable. To determine if centromeres are susceptible to adopt non-B-form DNA, we used a computational algorithm that models stress-induced structural transitions (SIST) for multiple non-canonical DNA secondary structures: Z-DNA, DNA melting (*i*.*e*. strand separation), and cruciform extrusions (Zhabinskaya et al., 2015). SIST was previously used by Kasinathan et al. to show higher probability to adopt non-B-form DNA for centromeres enriched in dyad symmetries (Kasinathan and Henikoff, 2017).

We ran segments of DNA in 5,000-bp blocks every 2,500-bp and took the maximum values for the overlapping regions whenever different. DNA transitions depend on temperature; since *D. melanogaster* is an ectotherm species, we ran SIST at five different temperatures at which *D. melanogaster* may be found (18°C, 22°C, 25°C, 30°C and 35°C) and determined enrichment probabilities for centromeres compared to their respective control regions. The probability of Z-DNA formation, which has not been previously analyzed for centromeres, is lower than controls for each of the centromeres irrespectively of the temperature and thus is unlikely to be associated with centromeres (**Fig. 2A**). As for cruciforms, only the centromere of the Y chromosome shows higher probability than controls at all temperatures (**Fig. 2B**). These findings are consistent with the observation that the Y is the only centromere showing an enrichment of inverted repeats (**Fig. 1F**), which are thought to adopt cruciform extrusions (Hamer and Thomas, 1974; Leach, 1994). Our findings in *Drosophila* are consistent with previous analyses on the centromeres of fission yeast, African green monkey and on human neocentromeres, where the probability of DNA melting was found to be higher than that of controls (Kasinathan and Henikoff, 2017).

**Figure 2:**
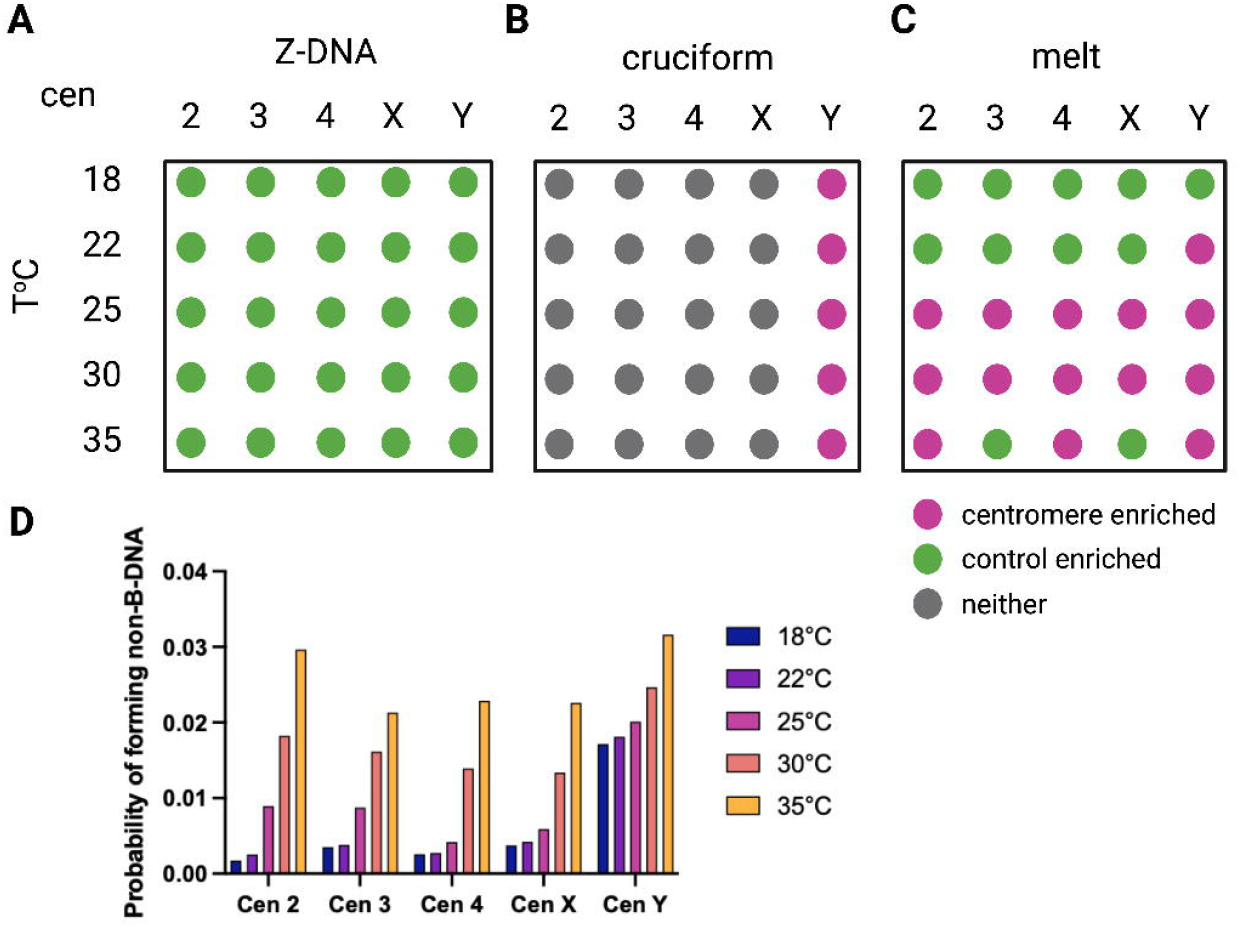
Enrichment of predicted non-B-form DNA at centromeres contigs using SIST. Diagram summarizing the SIST output. Results for Z-DNA (A), cruciform (B), and melt DNA (melt) (C) are shown for each of the centromeres at five different temperatures (°C). Different colors represent significance as outlined in the legend. (D) Average probability of non-B DNA formation for each centromere contig at different temperatures.

(Kasinathan and Henikoff, 2017). Interestingly, at 25°C and 30°C, all of the centromeres have higher probability than controls for DNA melting (melt). Centromere 2 and 4 display higher melting probability than controls also at 35°C. The Y displays higher DNA melting probability than controls at all temperatures greater than 22°C. At 18°C, none of the centromeres displays higher probability of DNA melting (**Fig. 2C**). When we plotted the overall probability of forming all three types of non-B DNA, we noticed that it increases with higher temperatures (**Fig. 2D**); this is likely due, at least in part, to the contribution of DNA melting to this probability. Cell and organism growth are regulated by temperature and the temperatures at which different organisms thrive are vastly different across eukaryotic species. If the ability of centromeres to adopt non-B DNA conformations needed for proper centromere function during cell division is also affected by temperature, this could be a factor under selection during evolution, contributing to the diversity of centromeric DNA sequences observed across lineages.

DNA melting is accurately predicted at actively transcribed regions that display strand separation *in vivo* (Zhabinskaya et al., 2015). As centromeres from across species have been shown to display low transcriptional activity (reviewed in (Mellone and Fachinetti, 2021)), the enrichment for this particular non-canonical DNA structure is especially interesting. DNA melting may facilitate transcription, which in turn could facilitate histone turnover or the formation of secondary DNA/RNA structures at centromeres, contributing to centromere specification (Kasinathan and Henikoff, 2017; Talbert and Henikoff, 2020).

### Enrichment of non-B-form DNA in centromeric contigs using GQuad

Previous work proposed that non-B-form DNA may be an evolutionary conserved signature required for centromere specification. Yet, aside from the Y centromere, which is enriched in inverted repeats and has higher probability of forming cruciforms than controls (**Fig. 1F** and **2B**), all other *D. melanogaster* centromeres show higher probability than controls only for DNA melting. As SIST only predicts 3 types of non canonical DNA structures, we wanted to expand our analysis to additional non-B-form DNA types. For this purpose, we used Gquad, a package that can predict 7 different non-B DNA structures: a-phased DNA repeats, G-quadruplexes, intramolecular triplexes (H-DNA), slipped DNA, short tandem repeats (STR,) triplex forming oligonucleotides (TFO), and Z-DNA. Gquad provides the positions and probability for specific non-B-form DNA using scores ranging from one asterisk (low likelihood) to three asterisks (high likelihood). In the absence of experimental data identifying non-B-form DNA and of a non-B-form DNA database for *D. melanogaster*, sequences known to form non-B-form DNA are not available as positive controls to determine the accuracy of our predictions. A previous study used inter-pulse duration (IPD) values (*i*.*e*. the time it takes to add a nucleotide during single-molecule sequencing) from PacBio long-read sequencing data to infer non-B-form DNA (Guiblet et al., 2018). When we plotted the average IPD values of regions predicted to form non-B-DNA (*e*.*g*. Z-DNA) identified by Gquad with a likeliness of two asterisks or greater in a 300-bp window centered on the sequence predicted to form Z-DNA, we observed IPD values that were twice as high, suggesting that the predictions generated by Gquad are accurate (**Fig. 3A**). Next, we calculated all the likelihoods for each type of non-B-DNA and combined them such that if a particular base pair was predicted to form non-B-form DNA of more than one type, the likeliness of the two were added together. To determine the significance of enrichment we used the two-sample Kolmogorov-Smirnov (KS) test. Through this analysis, we find that all centromeres are significantly enriched for non-B-DNA (**Fig. 3B-F**). Since the values for the 7 types of non-B-DNA are combined in this analysis, we next wanted to determine which types of non-B-DNA are contributing most to the enrichment of non-B form DNA at the centromeres found with Gquad. For this, we analyzed the enrichment of individual type and find that of the 7 non-canonical DNA forms, the ones that contribute the most are slipped DNA, STR, and G-quadruplexes (**Fig. 3G**).

**Figure 3.**
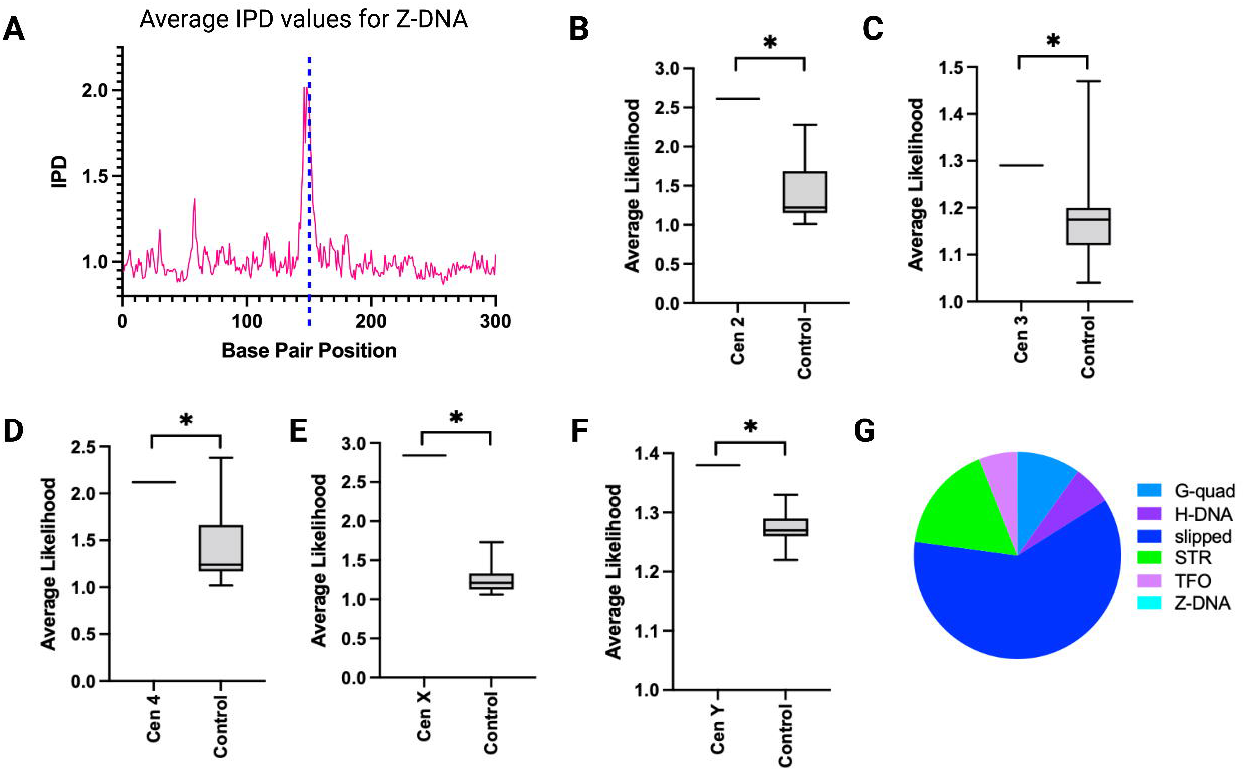
Enrichment of predicted non-B-form DNA in centromeric contigs using GQuad. (A) Plot showing the average IPD value for sequences predicted to form Z-DNA by GQuad with a likelihood of greater than two asterisks (see text for details). Z-DNA is centered around 150-bp. (B-F) Data distribution of likelihoods for each of the centromeres as a combination of all non-B DNA predicted by Gquad. Asterisks represent p<0.05 (KS test). (G) Pie chart showing the relative contributions of different non-B DNA types identified by Gquad.

Next, we sought to determine which types of repeats are contributing most to the likelihood of adopting non-canonical DNA secondary structures by ranking the average Gquad values for all repeats in the *D. melanogaster* genome. We find that simple satellite DNAs contribute the most, as they are consistently ranked higher than other elements (**Table S1**). Short satellites are known to be prone to form non-canonical DNA structures, particularly slipped DNA (Sinden et al., 2007). If centromeres need to be marked by unconventional DNA structures in order to function or be stable, a potential explanation for why satellite DNA is found at many regional centromeres across species could be that it can adopt non-B DNA.

To determine the prevalence of non-B-DNA at centromeric contigs compared to the rest of the genome (irrespective of GC content), we ranked all contigs that make up the genome based on the average Gquad likelihood. We find that all centromeric contigs fall within the top 37% of the 180 contigs picked up by Gquad as containing some form on non-B DNA, with centromeres X, 2 and 4 ranking 6th, 15th and 22nd, respectively (**Table S2**). These findings indicate that, although the centromeres may not rank the highest, they are among the most likely sequences in the genome to form non-B-DNA.

### G-quadruplexes are common features of *D. melanogaster* centromeres

To confirm our prediction of G-quadruplexes at the centromeres with an additional algorithm, we used G4Hunter, a more recent program that gives a G-quadruplex propensity score as output. Unlike Gquad, G4hunter takes into account G-richness and G-skewness of a given sequence. Furthermore, this algorithm was validated on published sequences known to form G-quadruplexes as well as with biophysical methods (Bedrat et al., 2016). We ran G4Hunter using a stringent threshold value of 1.5 and found that all centromeres, except the 3 and X centromeres, are enriched in G-quadruplexes compared to their respective controls (**Fig. 4A-E**). Having observed enrichment of G-quadruplexes with two independent methods, we conclude that G-quadruplexes are likely to be common features of *D. melanogaster* centromeres. G-quadruplexes play a role in transcriptional regulation, translation and replication (Bedrat et al., 2016). One possibility is that the higher prevalence of G-quadruplexes at the centromeres may contribute to centromere transcription homeostasis.

**Figure 4:**
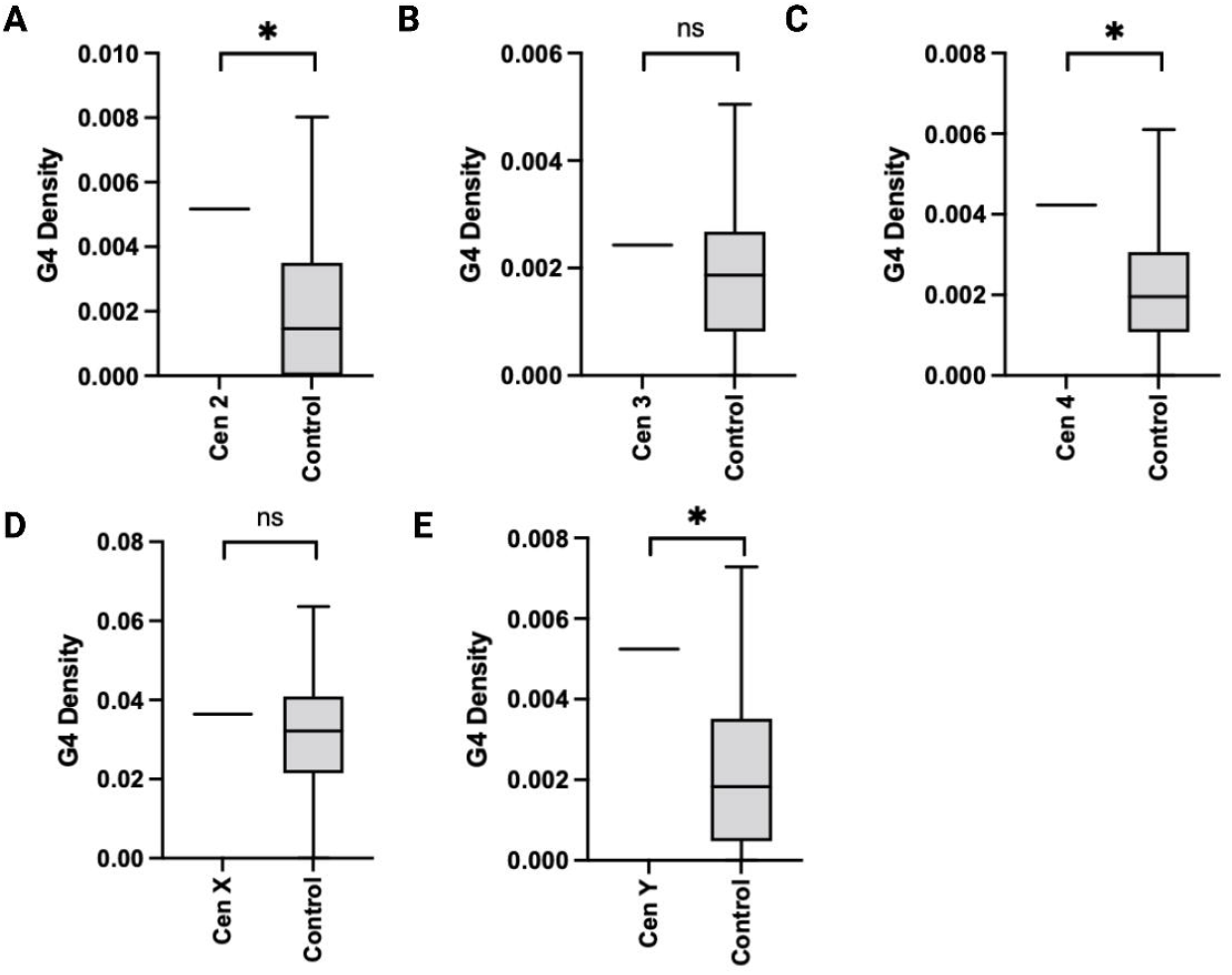
G-quadruplexes are common predicted features of *D. melanogaster* centromeres. (A-E) Graphs of the average G-quadruplex density for each centromere contig predicted by G4Hunter. Asterisks represent p <0.05 (One-sample t-test). Note that several control regions were not predicted to form any G-quadruplexes.

Collectively, our computational predictions suggest that *D. melanogaster* centromeres are enriched in non-B DNA secondary structures. Our findings are consistent with the model that non-canonical DNA forms may be evolutionarily conserved features of centromeres with possible functions in centromere specification. Under such paradigm, the only feature under selection at centromeres would be their secondary DNA structure. Since a myriad of primary DNA sequence combinations can accommodate such secondary conformations, such mechanism would enable ample opportunity for adaptation under intra-genomic conflict (Kasinathan and Henikoff, 2017).

## Acknowledgments

We wish to thank Patrick Grady, Sivakanthan Kasinathan, Asna Amjad, Marzia Cremona, Kateryna Makova, Amanda Larracuente, and Emmy Karim for discussions and suggestions and the UConn HPC and CBC for computing resources. This work was funded by National Institute of Health grant R35GM131868 to BGM and a Beckman Scholar fellowship to VSPP.

## Supplemental material

## Supplemental data

**Table S1. Table ranking the average Gquad value for all repeats in the *D. melanogaster* genome**. Repeats associated with centromere contigs are highlighted in yellow.

**Table S2. Table ranking all contigs that make up the genome based on the average Gquad likelihood**. Only contigs with an assigned likelihood are included (180 out of 190 total contigs in the genome). Centromeric contigs are highlighted in yellow

## Methods

### Genome data

The genome used in this paper is from Chang and Larracuente 2019 (Chang and Larracuente, 2019). The centromere contigs used for this analysis were Contig79 for centromere X, Contig119 for centromere 4, Y_Contig26 for centromere Y, Contig 3R_5 for centromere 3 and tig00057289 for centromere 2 (Chang et al., 2019).

### Source code

Code used to perform the analysis in this manuscript is available from GitHub (https://github.com/venkata14/dmel-nonb).

### Generation of controls regions

The controls used for the analysis were 50 random segments of the genome that are both the same size and have a similar GC content within 10% as the respective centromeric contig. A maximum of two controls with a 50,000-bp overlap was allowed.

**Detection of dyad symmetries using EMBOSS palindrome**

EMBOSS Palindrome (https://www.bioinformatics.nl/cgi-bin/emboss/help/palindrome) was used to detect dyad symmetries with the minimum palindrome being 5, the maximum palindrome being 100, allowing a gap limit of 20 and allowing overlapping dyad symmetries. We analyzed the output by calculating the dyad density, which we defined as the sum of the lengths of all palindromic regions identified by Palindrome divided by the length of the entire contig containing it. that contain that position. For a sequence, the length-normalized dyad density was defined as the sum of the values for each position divided by the sequence length.

### Prediction of Z-DNA, DNA melting and cruciform transitions using SIST

The probabilities of Z-DNA, Cruciform transitions and DNA melting were predicted using SIST (Zhabinskaya et al 2015) as described in Kasinathan et al. (Kasinathan and Henikoff, 2017). We used default parameters with the algorithm type “A” which uses the trans_compete C++ codes along with five different temperatures: 18°C, 22°C, 25°C, 30°C, 35°C for this analysis. For sequences greater than 10 kb in length, we slid a 5,000-bp window in 2,500-bp steps and analyzed these sub-sequences using SIST. The SIST predictions were then reassembled by taking the maximum SIST value for any given base pair.

To determine the the average probability of non-B-DNA formation for each temperature for all centromeres, we added the average value of Z-DNA, cruciform, and melt formation at each temperature.

### Prediction of non-B-DNA using Gquad

Gquad (v2.2-1; https://cran.r-project.org/web/packages/gquad/gquad.pdf) consists of multiple R packages that predict individual forms of non-B-DNA. We ran R packages on the heterochromatin-enriched *D. melanogaster* genome (Chang and Larracuente, 2019) for the 7 types of non-B-DNA: aphased DNA, G-quadruplexes, H-DNA, slipped DNA, Short Tandem Repeats (STR), Triplex Forming Oligonucleotides (TFO), and Z-DNA. The packages output likelihoods for each nucleotide from a range of one to three asterisks representing the likelihood of non-B-DNA formations. For those that did not output a likelihood, we used 2 asterisks as the default likelihood value. We then analyzed the data by combining all likelihoods for the 7 types of non-B-DNA for a respective sequence such that if there were overlaps in likelihoods of two different non-B-DNA types, we added those likelihoods together. This results in an array where each position is a summation of all likelihoods for a particular base pair.

### Identifying relative amounts of non-B-DNA using Gquad

Using the Gquad R package, we ran the package on the heterochromatin-enriched *D. melanogaster* genome (Chang and Larracuente, 2019) for the 7 types of non-B-DNA as similar to above. We then added all the positions predicted to form non-B-DNA for each of the 7 types and created a pie chart. To determine significance of prevalence between specific types of non-B-DNA in the centromere versus the controls, we used the one sample t-test on the average centromeric value and the control values for each respective non-B-DNA.

### Prediction of G-Quadraplexes using G4Hunter

G4Hunter (https://www.bioinformatics.nl/cgi-bin/emboss/help/palindrome) was run using a window size of 25 base pairs and threshold values of 1 and 1.5. The program outputs the positions of the nucleotides that are predicted to form G-Quadraplexes. Using these positions, we calculated the density of G-Quadraplexes by taking the total number of nucleotides predicted to form G-Quadraplexes and dividing them by the total number of nucleotides in the respective sequence.

### Validating non-B-DNA predictions of Gquad using IPDs

Publicly available PacBio sequencing reads from *D. melanogaster* (Kin et al 2014) were aligned to the heterochromatin-enriched *D. melanogaster* genome (Chang et al. 2019) with pbalign (SMRT v7.0), and IPDs were computed at nucleotide resolution with ipdSummary.py using the P5C3 chemistry (https://github.com/PacificBiosciences/kineticsTools/tree/master/kineticsTools). This outputs an IPD value which is an average of 3 IPD subheads values per nucleotide. All normalization of intermolecular variability and trimming for outliers was done automatically. Then, using the positive strand, all regions predicted to be Z-DNA by Gquad with a likelihood of two asterisks or higher were extracted in 300 base pair windows. The IPDs values of these sequences were extracted such that the predicted sequence to form Z-DNA was centered. All windows with no IPD values were filtered out, after which the IPD values of all sequences were averaged lengthwise and plotted.

### Statistical tests

The two-sample Kolmogorov–Smirnov test was used to compare distributions of SIST and GQuad likelihood values. One sample t-test was used for both the dyad density and G4Hunter distributions.

